# SAKE (Single-cell RNA-Seq Analysis and Klustering Evaluation) Identifies Markers of Resistance to Targeted BRAF Inhibitors in Melanoma Cell Populations

**DOI:** 10.1101/239319

**Authors:** Yu-Jui Ho, Naishitha Anaparthy, David Molik, Toby Aicher, Ami Patel, James Hicks, Molly Hammell

## Abstract

Single-cell RNA-Seq’s (scRNA-Seq) unprecedented cellular resolution at a genome wide scale enables us to address questions about cellular heterogeneity that are inaccessible using methods that average over bulk tissue extracts. However, scRNA-Seq datasets also present additional challenges such as high transcript dropout rates, stochastic transcription events, and complex population substructures. Here, we present SAKE (Single-cell RNA-Seq Analysis and Klustering Evaluation): a robust method for scRNA-Seq analysis that provides quantitative statistical metrics at each step of the scRNA-Seq analysis pipeline including metrics for: the determination of the number of clusters present, the likelihood that each cell belongs to a given cluster, and the association of each gene marker in determining cluster membership. Comparing SAKE to multiple single-cell analysis methods shows that most methods perform similarly across a wide range cellular contexts, with SAKE outperforming these methods in the case of large complex populations. We next applied the SAKE algorithms to identify drug-resistant cellular populations as human melanoma cells respond to targeted BRAF inhibitors. Single-cell RNA-Seq data from both the Fluidigm C1 and 10x Genomics platforms were analyzed with SAKE to dissect this problem at multiple scales. Data from both platforms indicate that BRAF inhibitor resistant cells can emerge from rare populations already present before drug application, with SAKE identifying both novel and known markers of resistance. In addition, we compare integrated genomic and transcriptomic markers to show that resistance can arise stochastically within multiple distinct clonal populations.

## Introduction

Compared to bulk RNA-Seq, where expression profiles are the result of averaging over millions of cells that may vary widely, single-cell RNA-Seq (scRNA-Seq) can be used to investigate the subtle but crucial differences in transcriptomic landscape that differentiate cellular state. Populations of cells that possess very similar gross cellular phenotypes might have remarkably different transcriptome profiles at the single-cell level due to stochastic transcription events, unsynchronized cell cycle stages, or inherent biological heterogeneity (Grün and van Oudenaarden 2015). Therefore, the normalization methods, statistical design models, and clustering methods of standard expression profiling may not work well for cell populations that cover wide ranges of cell types or conditions. One major concern in the analysis of scRNA-Seq datasets is the increased levels of noise in the measured transcript abundances per cell.

Excessive transcript dropout rates and stochastic bursting events in scRNA-Seq data create abundant non-detections, high variability, and complex expression distributions in the data as compared to bulk RNA-Seq. Therefore, it is important to distinguish low-quality, high-noise samples that are poorly amplified or degraded during library preparation. Optimal transcript normalization methods also differ for scRNA-Seq datasets as compared to typical bulk analyses. The mRNA content and capture efficiency can vary over a wide range between individual cells, the effects of which can be mitigated through the use of spike-in control molecules that can be used to model technical noise and as a function of transcript abundance (Brennecke et al. 2013; Ding et al. 2015).

Following normalization and quality control procedures, the next step in scRNA-Seq analysis often involves clustering of the samples to identify a set of gene markers that can segregate cells into distinct groups. Most published scRNA-Seq studies have used gene filtering and feature selection methods developed for bulk RNA-Seq, such as calculating the most variable genes (Klein et al. 2015; Macosko et al. 2015) the most significantly differentially expressed genes between known cell types (Shalek et al. 2013), or genes that have high contribution to the first few principal components (Satija et al. 2015; Li et al. 2016). This candidate set of marker genes is then used to identify subpopulations of cells via standard clustering methods, such as hierarchical clustering, k-means clustering, or principal component analysis (PCA). Visualization of these datasets using principal component analysis (PCA) or t-distributed stochastic neighbor embedding (t-SNE) (Maaten and Hinton 2008) can provide qualitative information about the number of clusters present and relative levels of cluster heterogeneity, but does not give a quantitative estimate of how many clusters are present nor whether a given sample belongs with one cluster or another. Proper choice of a clustering algorithm might depend upon the biological context of the samples. For example, most published single-cell clustering tools are optimized either for mixed populations of distinct cells (Kharchenko et al. 2014; Grün and van Oudenaarden 2015; Haghverdi et al. 2015; Satija et al. 2015; Xu and Su 2015; Zeisel et al. 2015b) or for time-series datasets that assume a smooth distribution from one cell type to another (Bendall et al. 2014; Marco et al. 2014; Trapnell et al. 2014; Setty et al. 2016). In practice, datasets can often include a mixture of cells from distinct cell types as well as related sub-clusters with significant overlap. Having a quantitative estimate of the relative similarity of each cell to each cluster centroid can be useful in these scenarios.

Here we present an integrated analysis tool that aims to facilitate the analysis of single-cell RNA-Seq data, addressing the challenges outlined above. Our Singlecell RNA-Seq Analysis and Klustering Evaluation (SAKE) method provides several modules that include: data pre-processing for quality control, sample clustering, t-SNE visualization of clusters, differential expression between clusters, and functional enrichment analysis. We applied SAKE to four recently published single-cell datasets with very different experimental designs (Deng et al. 2014; Ting et al. 2014; Zeisel et al. 2015a; Goolam et al. 2016) and evaluate its performance by the ability to correctly identify clusters reported in these studies. We also provide a comparison of SAKE performance on these datasets to multiple available scRNA-Seq analysis tools that have demonstrated rigorous performance in published studies: SINCERA, SEURAT, and SC3. We show that most scRNA-Seq analysis tools perform similarly for a wide range of sample types, despite each being algorithmically independent. In particular, SAKE performs similarly in most of these studies, and performs best for the Zeisel et al study that included a large, complex mixture of cells with extensive substructure. Importantly, SAKE also includes quantitative statistics to evaluate the clustering results, a feature missing from most other methods.

After demonstrating the success of the SAKE analysis package, we applied this method to better understand the mechanisms by which human melanoma cells respond to targeted inhibitors of the BRAF oncogene. Resistance to targeted BRAF inhibitors is widespread and presents a barrier to its efficacy as a therapeutic, since a large fraction of melanoma tumors initially respond to BRAF inhibition, but nearly all patients rapidly develop resistance (Müller et al. 2014; Shi et al. 2014; Sun et al. 2014; Perna et al. 2015). We used two single cell RNA-Seq platforms, Fluidigm C1 and 10x Genomics, to follow thousands of individual melanoma cells that have developed resistance to targeted BRAF inhibitors. Several published studies have identified validated gene markers of resistance to BRAF inhibitors, predominantly from bulk cell extracts, with often conflicting results that depend upon cellular context (Villanueva and Herlyn 2008; Villanueva et al. 2010; Konieczkowski et al. 2014; Sun et al. 2014; Ji et al. 2015; Verfaillie et al. 2015; Shaffer et al. 2017). We show that SAKE recapitulates several of these known markers as well as identifying novel markers of resistance that appear in rare populations of pre-resistant cells present in the population prior to drug application. In addition to these markers of “intrinsically resistant” cells, SAKE also identifies several genes that appear to mark a transient state between cells that are fully sensitive to BRAF inhibitors and fully resistant, as recently suggested in a study using single molecule FISH to evaluate resistance markers in melanoma cell populations (Shaffer et al. 2017). By integrating single-cell genomic and transcriptomic data, we show that this transiently resistant state can emerge from many clonal lineages present in the population prior to drug application. Together, these results suggest that many avenues of escape are available to melanoma cells responding to targeted inhibitors, and that the state of the cells prior to drug application can be an important determinant of which avenue is taken.

## Results

### Workflow for analyzing single-cell RNA-Seq data with SAKE

Single-cell RNA-Seq datasets contain both rich information as well as inherent technical artifacts. The SAKE workflow is designed to robustly categorize gene expression profiles while avoiding unwanted noise (Fig. 1A). Following the generation of a table of estimated gene abundance counts across samples; the next step in data analysis involves quality control steps to identify poorly amplified and problematic libraries. The SAKE workflow begins with this step. Samples with relatively low total transcript counts and gene coverage rates often represent degraded or poorly amplified libraries. These can be identified visually and removed from the sample set before proceeding with downstream analyses. The next step involves trimming the list of input genes to remove low abundance transcripts that suffer most from stochastic drop out events and technical noise issues. Median Absolute Deviation (MAD) is used as the preferred metric, but custom-filtering criteria can be implemented to filter out uninformative transcripts and those with expression levels below the reliable detection limit. SAKE provides a module to generate sample correlation heatmaps for use in evaluating the effects of filtering the gene list at a given MAD threshold or with additional criteria.

**Figure 1.**
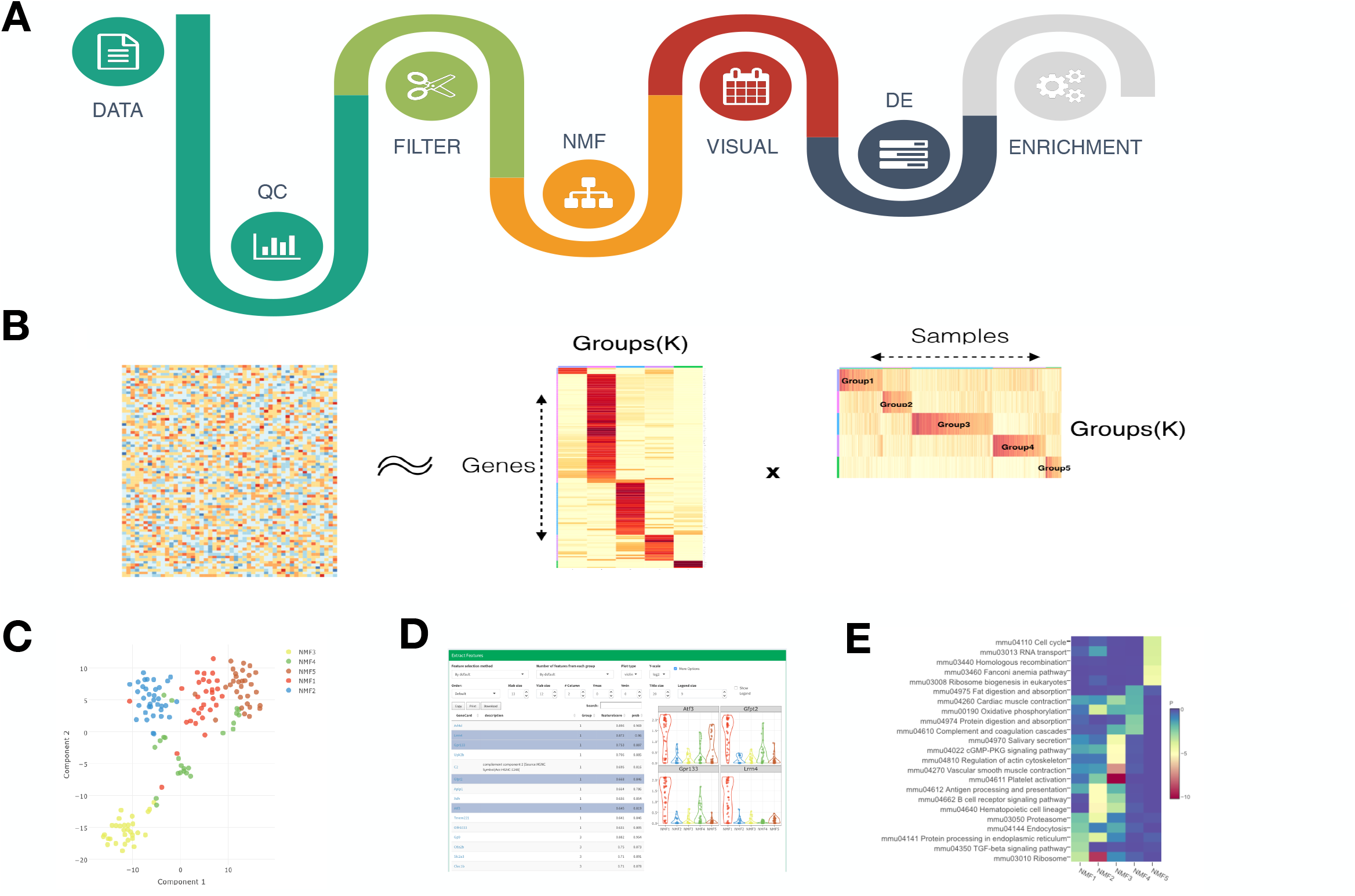
Flowchart of SAKE package and example analysis results. **A**) Analysis workflow for analyzing single-cell RNA-Seq data **B**) Schematic representation of the non-negative matrix factorization (NMF) method. A heat map of sample assignment and feature extraction from NMF runs, with dark red indicating high confidence in cluster assignments. **C**) A t-SNE plot to compare NMF assigned groups with **t-SNE** projections. **D**) A table of NMF identified features (genes defining each cluster) and a box plot of gene expression distributions across NMF assigned groups. **E**) Summary table for GO term enrichment analysis for each NMF assigned group.

The core of the SAKE clustering algorithm is a module to identify clusters via Non-negative Matrix Factorization (NMF) (Gao and Church 2005; Kim and Park 2007). NMF has successfully been used to identify molecular subtypes in bulk RNA-Seq expression profiles in many contexts (Hoadley et al. 2014; Weinstein et al. 2014; Moffitt et al. 2015; The Cancer Genome Atlas Network 2015; Yang and Michailidis 2016). Attributes that make NMF particularly appropriate for clustering of single-cell expression datasets include the ability to quantitatively estimate the number of clusters present in each dataset, *de novo,* and the ability to quantitatively estimate the likelihood that each sample belongs to a given cluster. Briefly, NMF attempts to factor a given gene expression matrix of N samples and M genes into two separate matrices: (1) a matrix of N samples belonging to k clusters and (2) a matrix containing the relative importance of each of the M genes in determining whether a sample belongs to each of the k clusters. This factorization can be attempted for a range of different values of k, with each iteration providing a quantitative measure of the robustness of cluster assignments upon randomization of the starting network. Information and update rules on how optimal matrix factorization is implemented is explained in extensive detail in the Supplemental Methods. To find the optimal value of k, one can minimize the residuals between the original full gene expression matrix (NxM) and the two factorized matrices (Nxk)(kxM) while simultaneously maximizing the cophenetic correlations between actual pairwise sample expression distances and the clustered dendrogram expression distances. SAKE provides a visual representation of matrix residuals and cophenetic correlation coefficients for a range of values of k to enable users select the optimal setting for the number of clusters, k, present in the dataset. Two quantitative metrics are given for choosing the optimal values of k: the silhouette index of the sample correlation matrix, and the cophenetic correlation coefficients.

Once an optimal number of clusters, k, has been determined, SAKE next performs a larger number of iterations of the NMF algorithm with fixed k, in order to robustly estimate the likelihood that each sample belongs to a given cluster and the relative importance of each marker gene in determining cluster membership. An example heat map of the cluster membership likelihood calculations for a particular N by k factorization of a gene expression matrix is shown in Fig. 1B (applied to the dataset of (Treutlein et al. 2016), described in detail in the supplemental section). In Fig. 1B, the full gene expression matrix is shown on the left, while the two factorized matrices as determined by SAKE are shown at the right. These results are presented as both a heatmap for easy visualization as well as formatted data table for quantitative comparisons. These quantitative estimates of cluster membership and gene marker importance are what give NMF an edge over more qualitative cluster evaluation metrics, such as t-SNE projection plots. Reassuringly, NMF identified clusters largely agree with t-SNE similarity maps, and the two metrics (NMF and t-SNE) are algorithmically independent; Fig. 1C presents a t-SNE plot of expression distances, colored by SAKE identified clusters in the dataset of (Treutlein et al. 2016).

Following cluster identification, SAKE provides several visualization options including: heat maps of gene expression markers enriched in each NMF cluster, t-SNE plots colored by NMF assigned cluster, and PCA plots colored by NMF assigned cluster. However, users can check the enrichment for additional marker genes of interest in each NMF cluster by alternately choosing to color each sample dot on these projection maps by the gene expression value of any chosen marker (Fig. 1D). Differential expression analysis between clusters can be evaluated using standard tools such as the DESeq2 algorithm (Love et al. 2014). Results are displayed for DE genes across NMF groups together with RefSeq annotation and Type-1 error corrected statistics (Fig. 1D). Moreover, GO Term enrichments and GSEA allow for the identification of functional categories enriched in each NMF cluster (Fig. 1E), which can serve as guidance for further investigation and follow up studies.

A detailed tutorial for using the SAKE analysis package is provided in the Online Methods and supplementary material. In addition to providing the code as an open source repository on Github, we present R markdown documents that allow for instructions, commands, and results to be presented together.

### Evaluation of SAKE on published data sets

We measured the success of the SAKE pipeline by its ability to reproduce the major findings from recently published scRNA-Seq studies that used a variety of analysis methods for cluster identification (Fig. 2A). The sample clustering results from the published scRNA-Seq studies served as the reference, and adjusted Rand index (ARI) and normalized mutual information (NMI) were used to evaluate the performance of SAKE as compared to a number of existing tools with results displayed for several recently published algorithms with demonstrated performance for scRNA-Seq analysis: SINCERA, SEURAT, and SC3. Fig. 2A presents a table summarizing the results for each of the three published algorithms on each of the four published datasets, together with the results from SAKE. Fig. 2B presents a table that explains the major differences between each of these three algorithms in terms of the methods used for gene feature selection and clustering. For comparison, we also include the results for a “simple” analysis that includes identifying the number of clusters, k, via a t-SNE projection plot, and then running a k-means clustering algorithm to assign samples to each of those k clusters (Fig. 2B).

**Figure 2.**
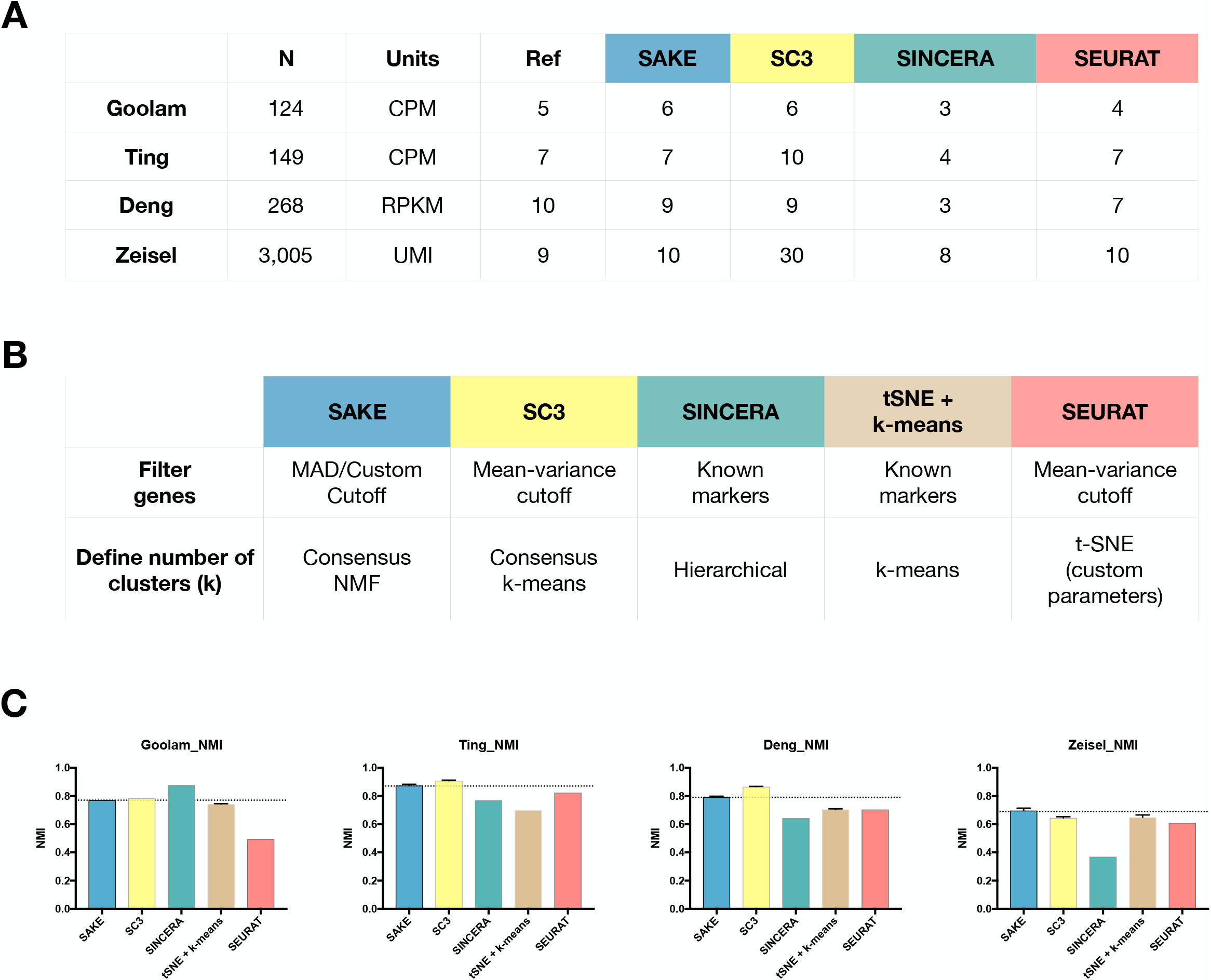
Data and performance summaries for scRNA-Seq software tools/pipelines. **A)** The number of samples and reported clusters from three published clustering methods (SC3, SINCERA, and SEURAT) as compared to SAKE (blue). **B**) Key features and techniques used by each method to perform gene filtering and to define the number of clusters. **C**) Normalized mutual Information (NMI) was used to compare the performance of each method on four published datasets in terms of the ability to recapitulate cluster assignments as given by the initial publication. Error bars were measured by subsampling 90% of the cells from each dataset and iterating 1000 times to ensure robust results.

To evaluate the robustness of these results, we randomly selected 90% of the samples for each of the published studies and performed 100 iterations of cluster identification for all of the compared clustering methods. In each iteration, we calculated the normalized mutual information (NMI) and adjusted rand index (ARI) using the cluster assignments in each of the published studies as the reference for accuracy. Results from these 100 runs are presented as bar charts in Fig. 2C that display mean NMI for each analysis method with error bars representing standard error. Supplemental Fig. S1 shows the results for ARI, another frequently used metric for evaluating accuracy of clustering results, which shows largely similar results, but can inflate small differences between differently sized clusters. Together these results demonstrate that SAKE is both accurate and robust across a wide range of sample sizes and experimental designs. Overall, each of these algorithmically independent clustering methods gives roughly similar results, with SAKE performance coming out best for experimental designs involving large complex substructures and large numbers of representative samples. While the ultimate assignments of each sample to a given cluster is largely similar for each of these algorithms, it is worth noting that SAKE is unique in providing quantitative metrics for the estimated number of clusters present, for the assignment of each sample to a given cluster, and for the relative ability of each gene to act as a marker for cluster membership.

### Application of SAKE to human Melanoma cell lines

Having demonstrated the success of the SAKE algorithm on published datasets, we next applied the SAKE method to questions that can best be answered by single cell analysis experiments: how cancer cells individually respond to targeted therapeutic agents. We first started with cells from a human melanoma cell line, *451Lu,* that carries an activating mutation in the BRAF oncogene. These cells have previously been demonstrated to be initially responsive to targeted BRAF inhibitor treatments (BRAFi), but to rapidly acquire resistance to these small molecule inhibitors (Villanueva et al. 2010). We defined cells in the naïve state before BRAFi treatment the “parental” population, *451Lu-Par.* By gradually increasing the dosage of BRAFi from 0.05um to 1μm on *451Lu-Par* and selecting for cells that survived after each round of treatment, we could derive a distinct population of BRAFi-resistant cells, *451Lu-BR,* that grew stably in a 1μm concentration of BRAFi (Fig. 3A). Differential response to BRAFi between the *451Lu-Par* and *451Lu-BR* cells was demonstrated through an MTT assay that measures metabolically active cells 72 hours following BRAFi treatment (Fig. 3B). Bulk RNA samples from these two conditions were collected and sent out for sequencing, and standard differential expression (DE) analysis was performed on these data. A handful of genes in the MAPK pathways were upregulated in *451Lu-BR* cells (Supplemental Fig. S2A), as well as 1000s of other genes of unknown functional relevance.

**Figure 3.**
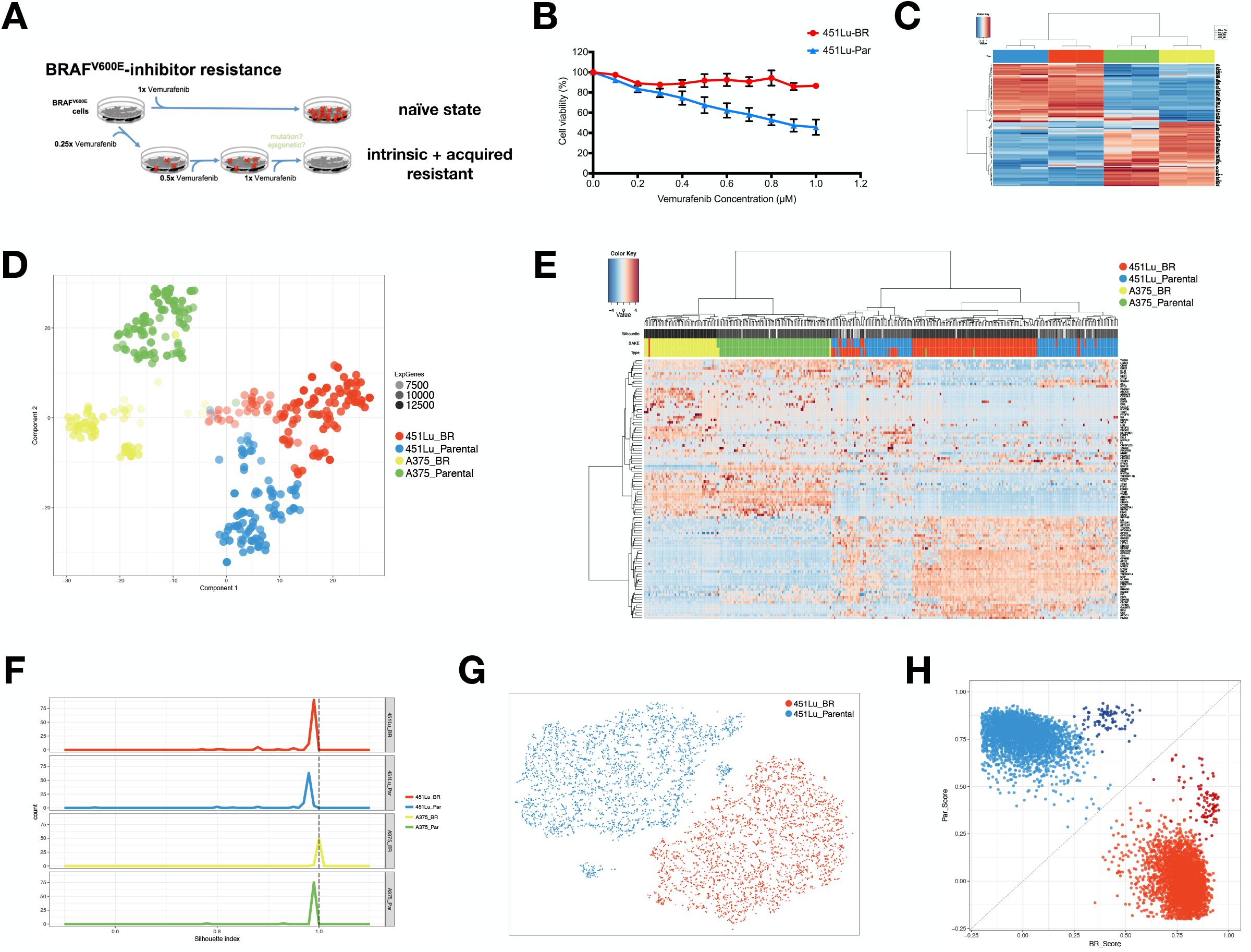
Bulk and single-cell RNA-Seq were used to study differential drug responses to BRAF inhibitor treatment. **A**) Naïve melanoma cells were treated with increased dosage of the BRAF inhibitor, vemurafenib, and cells that survived after each drug treatment were selected to gradually derive stably resistant BRAFi cell populations. **B**) Drug sensitivity was measured through the use of MTT assays to assess metabolically active cells 48 hours following BRAFi application. **C**) Melanoma signature gene sets were used to cluster bulk RNA-Seq data from melanoma cell lines. Samples first clustered by their cell types instead of the presence/absence of the drug treatment. **D**) A t-SNE map was used to displayed the expression profiles from ~400 parental and BRAFi resistant melanoma cells isolated using the Fluidigm C1 platform. The first t-SNE component separates the two cell lines, while the second component distinguishes between parental and resistant cells. **E**) Highly expressed and variable genes were used to classify Fluidigm C1 scRNA-Seq data. Higher levels of heterogeneity can be observed among ***451Lu*** cells as compared to ***A375*** cells. **F**) Distributions of silhouette index were used to assess cluster distances for each cell type. Lower silhouette index indicates a higher level of heterogeneity in cluster gene expression profiles, with 451Lu cell lines showing a lower average silhouette index in line with the heatmap shown in panel (f) **G**) To determine whether 451Lu cells have more intrinsic heterogeneity or more subclusters, 6545 scRNA-seq transcriptomes were obtained using the 10x Chromium platform. A t-SNE map of this 451Lu 10X data highlights two major groups of cells, corresponding to the parental and BRAFi resistant populations. **H**) In order to compare the 10x and C1 data on the same scale, a scoring system was implemented to score the distance of each cell from the population centroids of the parental (PAR) and BRAFi resistant (BR) populations. Spearman rank correlations to the centroid were then used classify each cellular transcriptome from both the C1 and 10X samples. All 451Lu cells from both the C1 and 10x datasets cluster together by cell type (Par/BR) with 10x cells showing slightly lower average scores due to sparse coverage.

To test whether these markers enriched in resistant populations can be generalized to other contexts, another human melanoma cell line with the same V600E mutation, *A375,* was selected and sent out for sequencing after deriving resistant populations following the same experimental procedure (Supplemental Fig. S2B, Supplemental Table S1). After combining data from *451Lu* and *A375*, melanoma signature gene sets (Hoek et al. 2008; Widmer et al. 2012; Verfaillie et al. 2015) were used as input for sample clustering (Fig. 3C). In general, the samples clustered first by their cell line instead of their treatment conditions. Differential gene expression analysis also showed that there was minimal overlap between the two cell lines in terms of shared genes that marked BRAFi resistant cells (Supplemental Fig. 2C), suggesting that these two cell lines that shared the same driver mutation might be using different pathways to enable growth in the presence of BRAF inhibitor molecules. One possible explanation is that these two cell lines possess unique expression profiles despite sharing the same genomic V600E BRAF driver mutation, and that these expression states contribute more to the overall transcriptome landscape than do the genes that drive resistance to BRAFi. To test that hypothesis, we would need to determine if our melanoma samples are derived from two distinct melanoma molecular subtypes, marked by their expression profiles, that generalize to additional samples.

Recent results from the Cancer Genome Atlas Consortium have described multiple transcriptome subtypes in melanoma patient tumors (The Cancer Genome Atlas Network 2015), similar to the “subtypes” that have previously been described for breast cancer and others (Hoadley et al. 2014). Indeed, like breast cancer subtypes, the melanoma cancer subtypes, dubbed “proliferative,” “invasive,” and “immune-infiltrated,” also correlate with overall patient survival rates. Moreover, these subtypes appear to be independent of driver mutations, since no enrichment was seen between the most common melanoma driver mutations (in BRAF, NRAS, and others) and transcriptome subtypes (The Cancer Genome Atlas Network 2015). We first used the SAKE algorithms, which work both for bulk and single-cell RNA-seq data, to determine whether we can reproduce these transcriptome subtype results on the TCGA datasets. Results from our SAKE analysis of TCGA data showed three groups with distinct biological functions and survival outcome that largely overlapped with the previously published findings. Supplemental Fig. S3 displays the SAKE clustering results with the original TCGA results in Supplemental Fig. S3A and the correlations with patient outcome in Supplemental Fig. S3B.

We next set out to determine whether these TCGA-derived melanoma subtype markers would also be present in cell culture samples, like the A375 and 451Lu cell lines described above. The markers of infiltrating immune cells were not expected in cell culture models, which was confirmed by very low expression of immune cell markers in the CCLE cell line data. Accordingly, the markers identified by SAKE in proliferative and invasive subtypes were used to classify a panel of Melanoma samples from the Cancer Cell Line Encyclopedia (CCLE). Indeed, *451Lu* and *A375* expression profiles were classified into the “proliferative” and “invasive” subtypes, respectively (Supplemental Fig. S3C). Each of the remaining CCLE cell lines also fell cleanly into one of these two groups. Moreover, BRAFi resistant cells from each of the *451Lu* and *A375* subtypes also remained within the same subtype as their respective parental cell lines, suggesting that subtype switching was not a dominant feature of BRAFi resistant cells (Supplemental Fig. S3C). Having confirmed that these melanoma transcriptome subtypes are present in both TCGA patient samples as well as the cultured cell lines described above, we next sought to determine whether these expression profiles would be faithfully represented by each individual cell in the population. Conversely, “subtype” might represent an averaging over many distinct cellular states that was not wholly present in any one cell. To answer this question, we would need to characterize the expression profiles of hundreds to thousands of melanoma cells from each subtype.

### SAKE Identifies Four Major groups in Fluidigm/Smart-Seq Datasets

Single-cell RNA-Seq data was prepared from both the *A375* and *451Lu* cell lines using the Fluidigm C1 system to isolate cells, convert mRNA to cDNA, amplify cDNA and single-cell libraries were generated using the Illumina Nextera XT kit. We isolated around 100 cells from each of the four conditions *(A375-Par, A375-BR, 451Lu-Par, and 451Lu-BR).* On average, more than 10,000 transcripts were detected per library (Supplemental Table S2). These C1 scRNA-Seq datasets were first mixed with the bulk data to evaluate the quality of the sequencing results in terms of transcript coverage. Signature gene sets that mark proliferative and invasive subtypes were used for clustering (Hoek et al. 2008). Most single cells recapitulated a similar expression pattern as that seen in the bulk RNA-Seq data and did not display a single-cell platform-specific profile (Supplemental Fig. S4A). Moreover, the similarity of the single-cell and bulk expression patterns suggest that the melanoma subtype profiles seen in bulk datasets are also representative of the dominant expression patterns of individual single cells. Additionally, the BRAFi resistant cells from both the *A375* and *451Lu* populations appear to remain within their molecular subtypes regardless of sensitivity to BRAFi treatment, as was seen for the bulk transcriptome profiles. Thus, response to BRAFi treatment may be constrained by molecular subtype, but the subtype markers cannot be used to explore BRAFi driven expression changes.

To explore the gene expression alterations associated with resistance to BRAFi, we next applied the SAKE algorithm to all ~400 cells from each of the *A375* and *451Lu* parental and resistance populations. A t-SNE projection of these results can be seen in Fig. 3D. The first component of the t-SNE projection separated out the two cell lines by their parental cell type *(A375* vs. *451Lu),* while the second component essentially separated the cells by their sensitivity to the BRAFi treatment, regardless of starting cell line. SAKE identified four different major clusters among the ~400 cells in the scRNA-Seq data (Supplemental Fig. 4B). The SAKE identified groups largely overlapped with the populations from which these cells derived (Fig. 3D), with little statistical evidence for isolated subclusters within each of the four populations. We did not observe a direct correlation between these four SAKE identified groups and cell cycle stages (Supplemental Fig. 4C). However, there were some deviations from these general trends. First, a small fraction of the cells in the parental cell lines appeared to exhibit markers of the resistant populations and, conversely, a small fraction of the cells isolated from the resistant populations appeared to still express parental markers as can be seen on the t-SNE plot of Fig. 3D and the heatmap of Fig. 3E. In addition, the t-SNE plot shows some sub-clustering within the *A375-BR* population, but these differences were not robust enough to represent a statistically robust independent population.

We next turned to ask whether any of the cell populations showed more or less heterogeneity in expression markers either before or after selection for resistance to BRAF inhibitors. Qualitatively, the heatmap of Fig. 3E shows that the parental and resistant *A375* cells separate cleanly by features identified using the SAKE algorithm. However, the parental and resistant *451Lu* cells appear to show more mixing in the cluster dendrogram, and more overlap in the expression of SAKE identified markers that distinguish parental and resistant cells. A quantitative analysis of the levels of expression heterogeneity in each population is displayed in Fig. 3F, where the SAKE confidence cluster assignment is plotted as a histogram for all cells in each of the four populations. While the *A375* cells were robustly placed in clusters with their cohorts, the *451Lu* cells show a larger range of confidence values, indicating a greater variance in the expression of cluster marker genes for these cells. One possible explanation is that the ~100 cells sequenced from each only sparsely sample each of several sub-clusters present in these cell groups. Another possible explanation is that *451Lu* cells are more intrinsically heterogeneous than the other groups sampled. Sampling a larger number of cells from these populations would enable the distinction between these possibilities.

### High throughput sparse 10x libraries for rare cell identification

To show that the results from the Fluidigm/Smart-Seq libraries were not singlecell RNA-Sequencing method-specific, we chose to sequence the same cell line using a different method. We selected the *451Lu* parental and resistant populations for higher throughput profiling on the Chromium Single Cell 3’ Solution from 10x Genomics, which provided information for many more cells with shallower coverage (Supplemental Table S2). Transcripts from ~6500 single cells from the *451Lu* parental and resistant populations were uniquely barcoded and sequenced, and PCR duplicates were removed using unique molecular identifiers (UMI). UMI counts for each cellular barcode were quantified and used to estimate the number of cells successfully captured and sequenced. On average, we obtained ~90,000 reads per cell and more than 5000 expressed genes in each cell (Supplemental Table S2). The Cell Ranger Single-Cell Software suite was used for demultiplexing, barcode processing, alignment, and initial clustering of the raw scRNA-Seq profiles. We first used t-SNE projection maps to get a general overview of the sequencing results, as displayed in Fig. 3G. In the *451Lu* parental cells, the majority of the cells fell into a single extended cluster, while two outlier groups formed unique subpopulations that were isolated and distinct from the rest of the parental population. These small isolated groups represented cells with extremely low or high transcript coverage rates, according to their UMIs, and were removed from further analysis. Cells from the *451Lu* resistant population formed a single group and did not have a clear separation into isolated sub-populations or distinct mixing with the parental population (Fig. 3G).

The sequencing depth and the number of expressed transcripts detected from the 10x data showed more variance as well as lower overall transcript coverage than the Fluidigm/Smart-Seq data (Supplemental Table S2), as expected from the lower total read counts sequenced per cell. To assess whether we could combine and compare these two types of data, we used the Nearest Shrunken Centroids method (Tibshirani et al. 2002) to calculate centroids from each population separately for parental and resistant cells, using only 10x-specific DE genes, assuming that genes detected in this shallow survey would be present in both Fluidigm C1 and 10x Genomics data (Fig. 3H). Briefly, the centroid of the expression patterns for each of the parental and resistant populations was calculated to obtain a median expression pattern for that population across all genes present in the 10x dataset. Spearman rank correlations were then used to score the distance of all cells in the sample from these centroids and this distance was the score for how “BR-like” or “Parental-like” each cell appeared (additional details are given in Supplemental Materials). Results from both the C1 and 10x data could then be displayed together (Fig 3H), and indicated that the cells largely separated according to their cell-type specific scores, and that most cells from each platform fell cleanly into one category or another. Moreover, the fraction of cells that showed equivalent scores for both parental and resistant profiles was similar across both the C1 and 10x platforms (Supplemental Methods). In addition, we compared the differentially expressed genes between the BRAFi resistant and parental cells for both C1 and 10x data and identified a largely similar set of commonly altered genes (Supplemental Fig. S5A-E), indicating that the two platforms give largely comparable results. As a result, the high degree of concordance between these two data sets shows that the findings correspond well to each other, with little platform-dependent effects beyond sequencing depth and the ability to identify rare cells in larger populations.

### DCT marks cells with intrinsic resistance in 451Lu melanoma cells

In the t-SNE project map of the C1 dataset (Fig. 3D), a small number of *451Lu-Par* cells can be seen located near the *451Lu-BR* population and distinct from the rest of the *451Lu-Par* populations (Fig. 3D-E). We wanted to test whether these rare parental cells that are exhibiting similar transcriptomic profiles to BRAFi resistant cells would also display less sensitivity to the BRAFi drug treatment. This would be consistent with an intrinsically resistant population already present in the parental cells.

We first used a standard differential expression analysis method, DESeq2, to determine the list of statistically enriched genes between the parental and resistant *451Lu* for C1 data, requiring two-fold mean expression changes and a p-value less than 0.05 (Fig. 4A). In addition, we used the default method that was implemented in the 10x Cell Ranger suite to identify differentially expressed genes in the 10x data. Checking through the list of genes that were identified as significantly differentially expressed by both DESeq2 and Cell Ranger, we saw highly variable levels of gene expression among these genes across resistant samples (Supplemental Fig. S5B-C). Therefore, we used SAKE to help identify genes more stably expressed in resistant populations from both Fluidigm C1 and 10x data sets. These SAKE markers have distinct expression distributions across parental and resistant populations, but highly stable expression within each population (Supplemental Figure S6A-B). Moreover, genes identified by both SAKE and DESeq2/Cell Ranger methods formed a high confidence candidate list for follow up validation studies (Supplemental Table S3).

**Figure 4.**
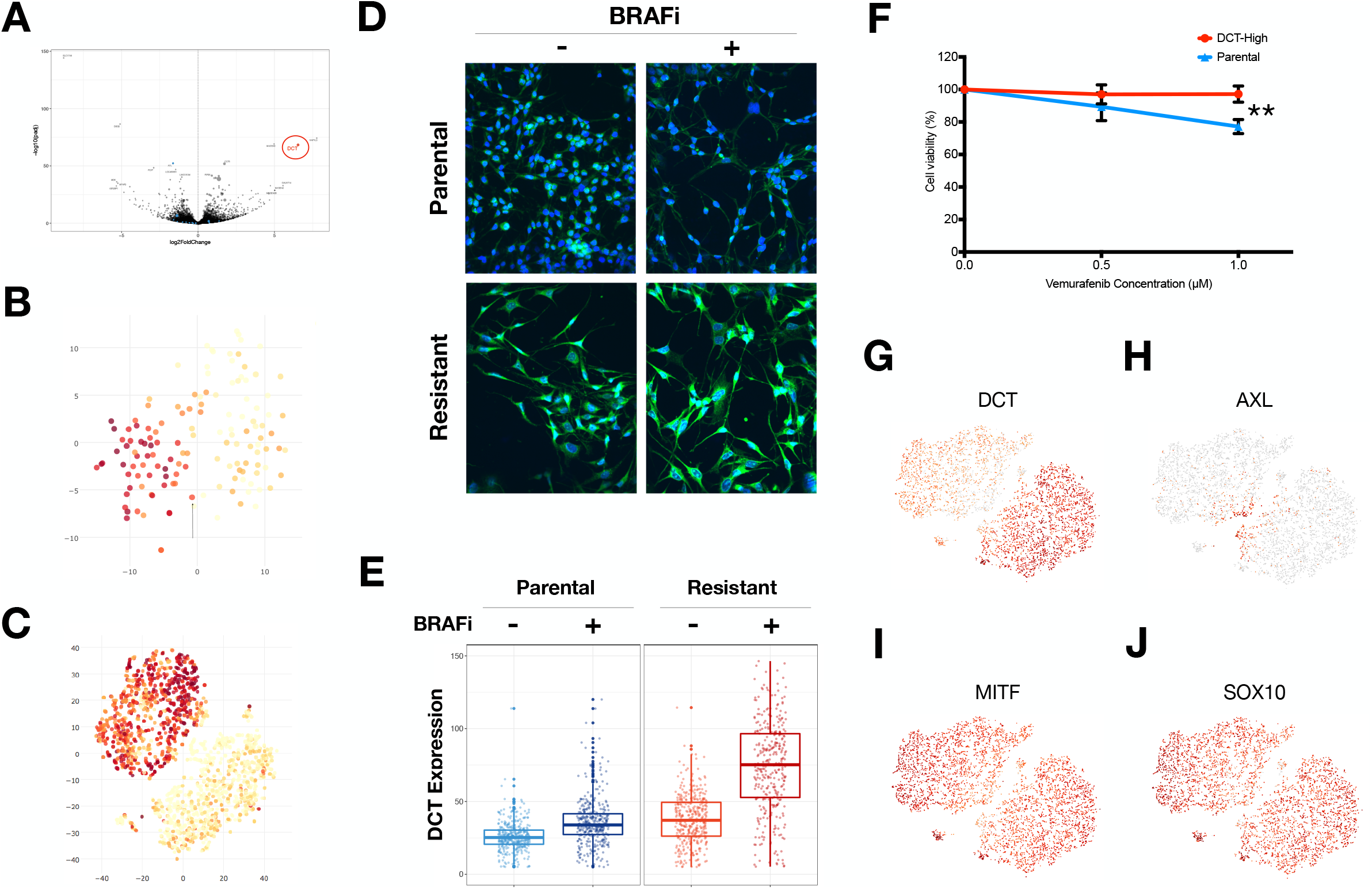
*DCT* marked *451Lu* melanoma cells that were intrinsically resistant to BRAF drug treatments. t-SNE maps were used to display the *DCT* expression levels from single-cell RNA-Seq data generated through **A**) Fluidigm C1 and **B**) 10X Genomics sequencing technologies. Most of the resistant cells had higher levels of *DCT* as compared to the parental cells. **C**) Volcano plot was used to display -log10(p-value) versus log2FoldChange between resistant and parental cells and to identify candidates for marking intrinsic resistant cells. A red circle highlights the DCT gene; blue dots highlight the potential candidate resistance markers reported by Shaffer et al. **D-E**) Cells were stained with a fluorescently labeled DCT antibody (green) showing that DCT also shows higher expression in BRAFi resistant cells at the protein level, quantified in the box plots of panel E. **F**) MTT assays measure metabolically active cells 48 hours after application of BRAFi to the media. DCT positive cells show significantly reduced response to BRAFi and higher survival rates. **G**) DCT shows binary expression pattern, with high levels in BRAFi resistant single-cells. **H-J**) Previously published markers of BRAFi resistance show undistinguished distributions in expression patterns across cells.

We then searched through these genes for cell surface markers that could be used to isolate cells. DCT was one of the highest confidence candidates that showed a distinct but consistent expression pattern between parental and resistant cells in both Fluidigm C1 and 10X datasets (Fig. 4B-C and Supplemental Fig. S5B-C). In addition, DCT is a membrane bound enzyme which makes it an ideal candidate for isolation by fluorescent based sorting methods. Parental cells with high DCT protein expression were isolated by FACS and tested for differential drug response following BRAF inhibitor challenge. These DCT-high parental cells showed a significantly reduced response to BRAF inhibitors, as measured by MTT assays for metabolically active cells 72 hours after plating the cells in media containing 1uM BRAFi compounds (Fig. 4F). This is consistent with DCT protein expression marking parental cells with reduced sensitivity to BRAF inhibitor treatment. To further confirm this, we stained *451Lu-Par* cells with DCT antibody and checked for the proportion of cells that survived after BRAF inhibitor treatment. If DCT-high parental cells show better tolerance to BRAF inhibition, there should be higher percentage of DCT positive cells remaining following application of BRAF inhibitor treatments to the naive parental population. Consistent with this, few DCT positive cells can be seen in the parental population prior to addition of BRAFi compounds to the media (Fig. 4D, left panels). Following treatment, the remaining cells can be seen to largely be DCT positive (Fig. 4D, right panels). As a positive control, most cells from the derived resistant *451Lu* population express high levels of DCT. These results are quantified and summarized in Fig. 4E.

We additionally checked a list of previously published genes that have been identified as associated with resistance to BRAF inhibitors. Surprisingly, most of these genes did not show a binary pattern similar to DCT, with dominant expression in one of the two populations (Fig. 4G-H). Instead, most of these published resistance marker genes showed a curious pattern of expression that was most differential at the interface between the two populations (Supplemental Fig. S7). One possible explanation for this expression pattern would present a model where expression of some genes is required to initiate the process of developing drug resistance, but this expression is not required for maintenance of the drug-resistant state. The level of support for this model is explored next.

### Support for a transient transcriptional state allowing for acquired drug resistance

A recently published study reported the use of bulk RNA-Seq and high throughput single-cell FISH techniques to identify a panel of signature genes associated with BRAF inhibitor resistance in human melanoma cells (Shaffer et al. 2017). This study proposed that selected genes identified previously in the literature, such as WNT5A, AXL, EGFR, PDGFRB, and JUN could mark a transitional stage in which cells are better able to develop resistance to BRAFi treatment, a state they dubbed “pre-resistant.” More interesting, after removal of BRAF inhibitors, these intermediate cells were more likely to revert to their naïve parental state and become drug sensitive again. Intrigued by this finding, we checked the expression distributions for a subset of these transient markers in the 10x Genomics data (Supplemental Fig. S7). While we did not see enrichment for these markers in any isolated outlier groups, these markers were enriched at the tip of the parental population that was proximal to the resistant populations as well as at the tip of the resistant population that was proximal to the parental cluster. These results would be consistent with a transient population between the two fully drug sensitive and fully resistant states.

We then ran SAKE on these 10x data to determine whether SAKE would also identify the presence of this unique “cluster” of candidate transient cells based solely on their transcriptional profile without any *prior* knowledge from the marker lists (Supplemental Fig. S7). SAKE reported 3 major clusters from the entire 10x dataset, which were the parental cells, resistant cells, and a new population of cells that sat at the interface between these two major clusters on the t-SNE projection maps (Fig. 5A and Supplemental Fig S8A). These (yellow) intermediate cells could represent parental cells existing in a “pre-resistant” state as well as cells in the drug-resistant population that are exiting this transient state, consistent with the scenario proposed in Shaffer et al., (2017). We next used the SAKE identified markers of this intermediate population to determine whether the genes that marked this population showed overlap with previously identified resistance-associated gene sets.

**Figure 5.**
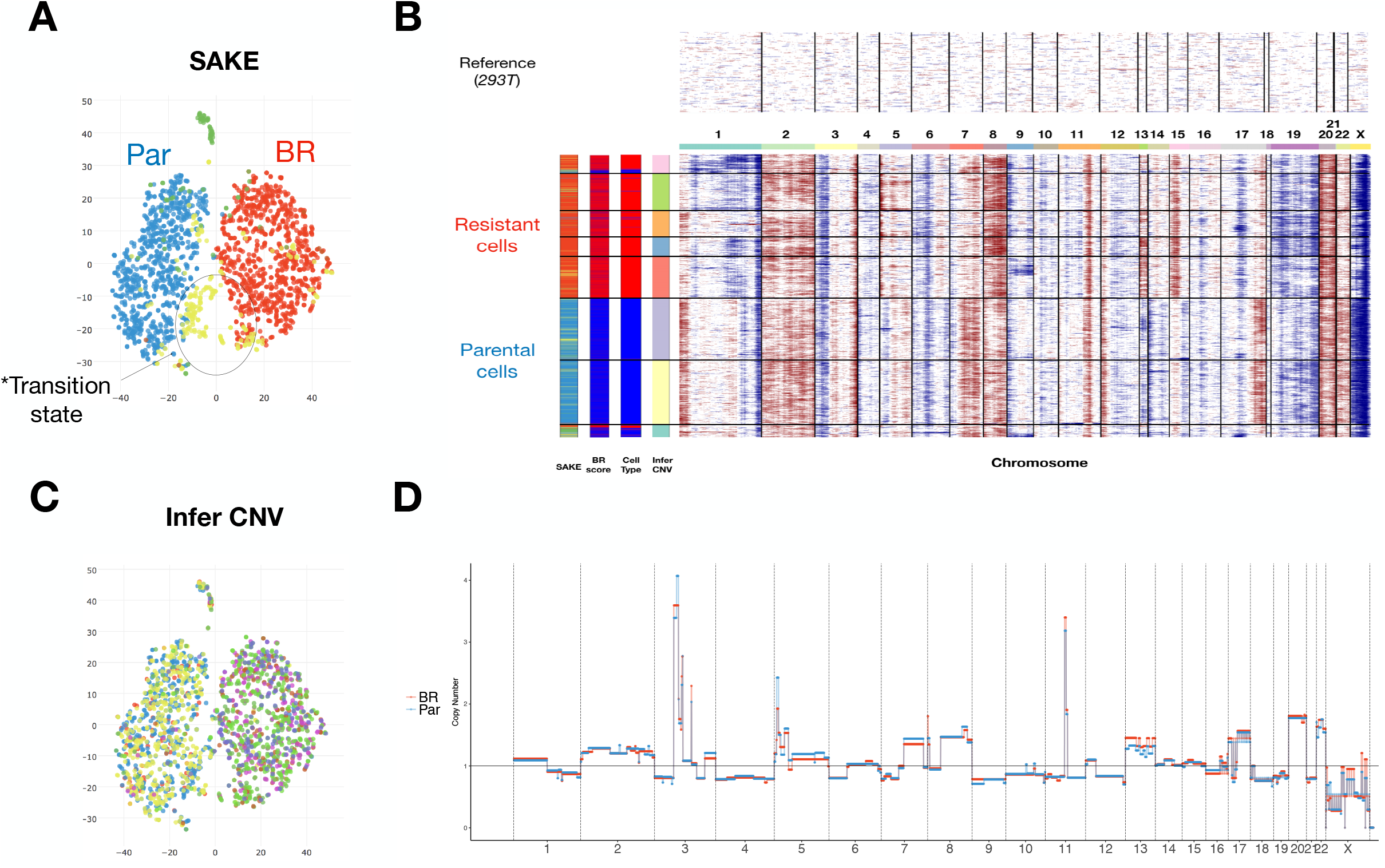
CNV and transcriptomic profiles identified an intermediate state between parental and resistant cells. **A**) t-SNE map was used to display the single-cell RNA-Seq data from melanoma single cells. Each circle represents a cell and is colored by SAKE identified groups. One group of cells (yellow) marked the potential transitional state during the acquisition o1 BRAF drug resistance. **B**) Inferred CNV profiles (scRNA-Seq) were used as a proxy to assess the clonal states among parental and resistant cells. **C**) t-SNE map was used to display the clonality among single-cells. Each circle represents a cell and is colored by inferred CNV groups. No clear enrichment for particular clonal population can be found for cells in the transitiona state. **D**) Copy number profiles from bulk DNA samples were used to validate the patterns observed from inferred CNV profiles (scRNA-Seq).

Consistent with the previously reported melanoma BRAFi resistance associated genes, the intermediate populations were marked by high levels of EGFR, AXL, JUN, WNT5A, FGFR1 and NRG1 (Supplemental Fig. S8C-D). These cells also expressed the lowest levels of differentiated melanocyte lineage genes such as MITF (Supplemental Fig. S8B) and SOX10. These findings are consistent with the expression patterns displayed by the transient population previously identified by Shaffer et al., (2017); this supports the hypothesis that some melanoma cells acquire resistance by transiting through a transient “pre-resistant” state marked by high levels of MAPK pathway genes such as EGFR and AXL, but that the expression of these genes is not required for the maintenance of a stably resistant population. In addition, SAKE identified several additional genes that mark this candidate “transitional” population, which are given in Supplemental Table S4. While these cells do not occupy an isolated sub-cluster on the t-SNE projection map, the SAKE algorithm did identify this population as a robust subcluster, highlighting the success of SAKE at finding small sub-populations in high dimensional data.

To garner additional support for the hypothesis that these cells represent a transient transcriptional state that can seed the resistant population, we also generated genomic copy number data to compare to the transcriptome data. Integrating both RNA expression data and genomic CNV data would allow us to determine whether particular subsets of the population with similar expression patterns were also more likely to share CNV mutational patterns, as might be expected through simple selection for genetic mutations. Conversely, if many cells with highly divergent CNV mutational patterns nonetheless shared very similar transcriptional patterns, one would expect that the “transitional” expression state might be due to additional factors that could reflect epigenetic or gene regulatory modifications. This is conceptually similar to the Luria-Delbrück fluctuation analysis that was performed in the Shaffer et al., (2017) study to demonstrate that resistance to BRAFi treatment in their cells could be due to non-heritable effects arising stochastically among cells from several clonal populations (Shaffer et al. 2017). If random genetic mutations were giving rise to resistant cells in this scenario, one would expect that a small number of cells would spontaneously develop resistance, and that these cells may display distinct genomic mutations patterns while potentially displaying shared transcriptomic patterns.

To check whether this phenomenon holds true in our *451Lu* melanoma data, we first performed copy number variation (CNV) analysis from the scRNA-Seq data, as has been described previously (Patel et al. 2014; Tirosh et al. 2016). Briefly, genes were binned into sliding windows containing 100 consecutive genes, to ensure that the alterations due to differential expression patterns would be averaged out over a large enough region such that very large CNVs could be detected independent of expression differences. In addition, 100 karyotypically normal *Human 293T* cells with scRNA-Seq profiles of approximately the same depth were used to derive the baseline reference for calibrating relative CNV calls and to validate that this method would not incorrectly call CNV patterns in karyotypically normal data (Fig. 5B). To ensure that the CNV profiles we inferred using this method match what would have been found as directly measured from the DNA, we additionally isolated DNA from bulk cell populations from each of the parental and resistant populations and generated DNA libraries directly (Supplemental Table S4). We found a high degree of concordance between the CNV profiles derived from the scRNA-Seq data and the bulk DNA-Seq data (Fig. 5B,D). This suggested that the thousands of cells sequenced using the scRNA-Seq protocol could also be used as proxy to infer clonal CNV relationships.

The inferred CNV patterns showed two major clones that corresponded to the parental and resistant cell populations. Depletions on chr9p and amplifications on chr1p, chr7p, chr11q, chr20q, and chr22q were signatures for the resistant cells, and these large regions of genomic alterations match what has been reported in other CNV profiles using bulk melanoma samples (Beroukhim et al. 2010). Beyond the two dominant clones, these cells were further classified into 8 subgroups, or “clonal lineages.” However, none of these 8 subgroups of cells showed an enrichment for the cells that are identified as having the “transitional” transcriptome patterns (yellow bars in Fig. 5B), which could be found in several of the 8 subgroups. This suggested that these cells were derived from several unrelated clonal lineages, and were likely not derived from a single strictly heritable lineage. This is consistent with the hypothesis that non-genetic effects contribute to the acquisition of resistance to BRAF inhibitors in melanoma cells, and is consistent with previous reports (Shaffer et al. 2017; Sharma et al. 2017).

## Discussion

Methods developed for bulk RNA-Seq data analysis (Anders and Huber 2010; Robinson et al. 2010; Dillies et al. 2013; Love et al. 2014) can be adapted for application to scRNA-Seq data, but need to be tailored to address the specific issues inherent in working with noisy, low-coverage, heterogeneous sample sets. We present a processing and analysis pipeline for single cell RNA-Seq datasets, SAKE, that is designed to look for quality control issues specific to scRNA-Seq data, identify sub-clusters present in the cell populations, and evaluate the gene and functional group enrichments in each cluster. Importantly, SAKE provides quantitative estimates of cluster membership alongside qualitative evaluations of cluster relationships via PCA plots and t-SNE projection maps. We show that SAKE can provide very similar single-cell cluster results to those derived from a variety of sample sources and evaluated with very different algorithms. We also show that SAKE provides accurate and robust results for a wide range of experimental designs. Moreover, SAKE operates as a simple, intuitive pipeline where the parameters are set by quantitative evaluation criteria provided by SAKE during run time, and quantitative metrics for the clustering results are presented as output.

Having verified the success of SAKE on several published scRNA-Seq datasets, we next applied this method to uncover the transcriptional alterations that occur as melanoma cells develop resistance to targeted inhibitors of the BRAF oncogene. Single cell transcriptomes were obtained from both a low-throughput high-depth method (Fluidigm C1, ~100 cells per group) as well as a high-throughput shallow sequencing technology (10x genomics, ~3500 cells per group). The expression profiles and typical melanoma specific marker levels were very similar between these two datasets. Overall, these two sequencing technologies provided highly concordant results, suggesting that platform-specific effects did not dominate the results.

### Identification of rare “intrinsically resistant cells”

The datasets from the Fluidigm C1 and 10x platforms were combined to identify statistically differentially expressed genes between the *451Lu* parental and resistant populations. Those genes that were highly abundant in the *451Lu* resistant cells, with much lower overall expression in the *451Lu* parental population, formed a candidate list of resistance marker genes. We next sought to determine whether rare cells in the parental population might be expressing these candidate resistance markers. We validated one novel resistance marker gene, DCT, with high expression in the majority of the BRAFi resistant cells and low expression in 99% of the parental population. Using a fluorescent antibody for DCT, these rare DCT positive cells were sorted from the parental population and challenged to determine sensitivity to BRAFi treatment. Consistent with DCT marking cells that are more likely to be resistant to BRAFi treatment, the DCT positive population showed reduced sensitivity to BRAFi challenge. While one report identified DCT as a potential marker for general resistance to radiotherapy (Pak et al. 2004), DCT has not previously been evaluated for its role in resistance to targeted inhibitors or specifically BRAF inhibitors.

### Cell type specific markers for conferring resistance

Surprisingly, most of the genes that have been identified in previous studies to confer resistance to BRAFi treatment in melanoma did not show a similar expression pattern to DCT. That is, most of these genes did not show an overall binary distribution with high levels in most resistant cells and low expression in most parental cells or vice versa. Instead, a large fraction of these genes showed differential expression just at the boundary between the parental and resistant populations. This expression pattern could be consistent with previously proposed models that suggest a transitional cellular state dubbed “pre-resistance” in which cells transiently express these markers as they develop resistance to BRAF inhibitors (Shaffer et al. 2017; Sharma et al. 2017), but do not necessarily continue marker expression once resistance is stably established. In this model, one might expect cells with shared transcriptomic patterns, marking the transitional state, to not necessarily share genomic mutational patterns. While previous studies have used Luria-Delbruck analysis to infer the presence of multiple clones with non-heritable resistance patterns (Shaffer et al. 2017), having direct evidence for distinct mutational patterns among cells with shared transcriptomes would extend support for this model.

### Integrating genomic and transcriptomic data to infer lineage

Integrating genomic mutational information allows for an explanation of the clonal relationship of the cells that were selected for resistance to BRAF inhibitors. If most cells in the resistant population appear to be more genetically similar than the cells in the parental population, this would imply a bottleneck scenario where a small number of clones were selected. On the other hand, if resistance arises stochastically in many genetically distinct clones, this would imply that non-genetic mechanisms could be contributing to the development of drug resistance. Our data seem to support a contribution from both mechanisms. Specifically, the intrinsically resistant DCT positive cells likely mark a small number of clonally related cells that were innately resistant to BRAFi treatment prior to drug application, and which were simply selected from the population. However, most of the previously published melanoma markers appear to be expressed in a small number of cells from both the parental and resistant populations, and these cells seem to cluster in a similar expression space on the t-SNE projection maps. This particular pattern could be consistent with a model in which clonally unrelated cells stochastically develop expression patterns that allow them to survive in the presence of inhibitors. This model would imply a non-heritable mechanism for acquired resistance that would derive from multiple clonal lineages rather than a small number of selected clones. Integrating genomic CNV and RNA-Seq information showed that the candidate “transitional” cells appear to derive from many distinct clonal lineages, and showed no enrichment for any particular clonal lineage. Moreover, cells with very distinct expression patterns were also present within the same clonal lineage, suggesting that the transitional expression pattern was not clonally inherited. These results highlight the complexity of melanoma cell populations and the multiple mechanisms available to respond to targeted inhibitor therapies.

## Materials and Methods

### Cell Culture

Parental cell lines A375 and 451Lu were cultured in DMEM media supplemented with 10% FBS and 1% Pencillin-Streptavadin. Resistant cell lines A375-Br15 and 451Lu-Br3 were cultured in DMEM media supplemented with 10% FBS, 1% Pencillin-Streptavadin and 1uM of BRAF inhibitor, PLX4720. PLX4720 was obtained from Selleckchem (Catalog No.S1152).

### RNA-Seq library preparation for bulk melanoma cell samples

Total RNA was extracted from approximately 10^Λ^6 freshly collected melanoma cells following standard Trizol RNA extraction protocols. RNA-seq libraries were prepared from 500ng of total RNA using the Illumina Tru-seq Stranded Total RNA kit. Libraries were barcoded and pooled in order to obtain a minimum of ~40 million reads per library on the Illumina HiSeq 5000 platform.

### RNA-Seq library preparation using the Fluidigm/Smart-Seq platform

RNA-Seq library preparation Cultured cells (70-80% confluency) were trypsinized and resuspended into a single cell suspension in media. Cells were then diluted to 175-300 cells/ul and loaded onto a medium (10-17 uM) Fluidigm C1 IFC. Single cell capture was done on the IFC and the number of cells captured at each site was noted using a phase contrast microscope. Only single cells were used for analysis. cDNA was amplified on the chip using the SMART-Seq v4 Ultra Low Input RNA Kit for Sequencing (Clontech) and quantified using picogreen assay. ERCC (External RNA Controls Consortium) RNA spike-in Mix (Ambion, Life Technologies) was also added to the reaction. Libraries for sequencing were prepared in 96 well plates using the Illumina Nextera XT DNA Sample Preparation kit according to the protocol supplied by Fluidigm. Final quantification by bioanalyzer using Agilent’s High Sensitivity DNA Analysis Kit as well as qPCR. Libraries were then sequenced 76bp single end on Illumina NextSeq platform to a depth of 5-15 million reads per cell.

### RNA-Seq library preparation using the 10x Genomics platform

Samples were submitted to Genome Technology Center at NYU Langone Health for processing on 10x Genomics Chromium and sequencing. A Chromium Single Cell 3’ Library and Gel Bead Kit V2 (PN-120237), Chromium Single Cell 3’ Chip Kit V2 (PN-120236) and Chromium i7 Multiplex Kit (PN-120262) were used with a 10x Genomics Chromium for Single-Cell Library Preparation Instrument, per the manufacturer’s specifications and manuals and then sequenced paired-end 150 bp on HiSeq 4000 to a depth of 90000 UMI per cell.

### Bulk DNA-Seq library preparation

Cultured cells at 70-80% confluency were trypsinized and counted. DNA was extracted from 3-4 million cells using Qiagen DNeasy Blood and Tissue kit following the manufacturer’s instructions. DNA output was measured using a nanaodrop. 2ug DNA was fragmented using a Covaris sonicator. Following this, libraries for sequencing were prepared by subjecting the resulting DNA fragments to end-repair, 3’ adenylation and ligation of TruSeq barcoded adapters. After ligation, DNA fragments were size selected using Ampure XP beads. A 0.75 ratio of beads to sample was used to select fragments larger than 200bp and avoid adapter dimers. These fragments were further amplified by PCR, cleaned using Qiaquick PCR Purification Kit and then once again size selected using Ampure XP beads with a 0.75 ratio of beads to sample. The final libraries were then quantified on Nanodrop and Bioanalyzer using Agilent’s High Sensitivity DNA Analysis Kit. 10nM of each sample was then pooled together and the resulting pool was then sequenced single read 76bp length on the Illumina NextSeq to a depth of 6-7 million reads per sample.

### MTT Assays for measuring cell response to BRAF inhibitors

Parental (A375, 451Lu) and Resistant (A375-BR, 451Lu-Br3) cells were plated in 96 well plates at a concentration of 7500-10000 cells/well and allowed to culture overnight. Once the cells were at 70-75% confluency, cells were treated in triplicates with different concentrations (0.1uM - 1.0uM) of PLX-4270 in 200μl of media for 72 hours. After the treatment, the media with drugs was removed and replaced with 100ul fresh phenol red free media without the drug. MTT Assay was then carried using Vybrant^®^ MTT Cell Proliferation Assay Kit from Thermo Fisher Scientific.

### FACS sorting to isolate populations expressing identified markers

Cells were trypsinized and resuspended in FACS buffer (PBS with 0.1% BSA and 0.05% Sodium Azide) and stained with Anti-DCT antibody conjugated to Alexa Fluor 488 (TRP2 Antibody (C-9): sc-74439 from Santa Cruz Biotechnology). Stained cells were then run on Becton Dickinson FACS Aria (SORP) Cell Sorter and cells expressing High DCT were bulk sorted into DMEM media supplemented with 10% FBS and 1% Pencillin-Streptavadin. The sorted cells were then immediately plated on 96-well plates at a concentration of 5000 cells/well. Cells were allowed to grow to 70-75% confluency before PLX-4720 treatment. PLX-4720 treatment (0.5uM and 1.0uM) was done for 72 hours and then MTT assay was performed using Vybrant^®^ MTT Cell Proliferation Assay Kit from Thermo Fisher Scientific.

### Imaging

Cells were plated on glass coverslips coated with Poly L-Lysine solution (0.1% w/v from Sigma Aldrich) and allowed to grow for 24 hours before drug treatment. Cells were treated with either 0.5μM or 1.0μM concentration of PLX-4720 for 2 hours, 24 hours, 48 hours and 72 hours. Cells after treatment were washed with PBS and fixed using ice-cold Methanol. Fixed cells were blocked with 5% FBS in PBS overnight at 4°C and stained with a 1:50 concentration of Anti-DCT antibody conjugated to Alexa Fluor 488, diluted in 5% FBS in PBS overnight at 4°C. This was followed by NucBlue^®^ Fixed Cell ReadyProbes^®^ Reagent (Cat No R37606, Thermo Fisher) counterstaining of nuclei according to the manufacturer’s manual and mounted on a slide using ProLong™ Diamond Antifade Mountant. The imaging was done on a confocal microscope

### RNA-Seq data processing

STAR 2.5.2b (Dobin et al, 2012) was used to align sequencing data to the Human hg19 reference genome with gene transfer format (GTF) file downloaded from UCSC (on date 2016-05-25). For bulk and Fluidigm C1 single-cell RNA-Seq libraries, samples with less than 80% uniquely mapped reads were filtered out before downstream analyses. Kallisto 0.42.5 was then used for abundance estimation using the same GTF file to get the gene expression matrix, M. Log-transformation is then applied on this gene expression matrix, as M’ = log2(M + 1). DESeq2 was used to identify differential expressed genes between conditions.

### SAKE clustering

SAKE works on gene expression matrix, ***M***, as input. Columns correspond to cells/samples and rows correspond to genes/transcripts. Each element of ***M*** corresponds to the expression of a gene/transcript in a given cell. By default, SAKE does not perform any normalizations or corrections for cell cycle or batch effects. The users should perform gene/transcript abundance estimation before feeding the gene expression matrix into SAKE for clustering, feature extraction, and other downstream analysis.

### Normalized mutual information

For the published datasets we chose to evaluate the performance of clustering tools, the normalized mutual information (NMI) is calculated to compare the similarity between the SAKE (and other tools) clustering results and the published cell-labels. To calculate NMI, we first construct confusion matrix and derive accuracy and precision from it. By counting the number of overlapping points for each pair of clusters between *C(r)* and *C(g),* we can construct a summary table. The mutual information can be calculated via:

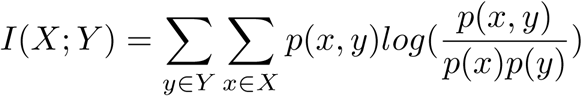

where *p(x,y)* is the joint probability distribution function of X and Y, and *p(x)* and *p(y)* are the marginal probability distribution functions of X and Y respectively. The normalized mutual information can then be calculated by accounting for the entropy in each of X and Y. First, entropy of a cluster **w**

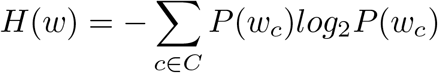

where: **c** is a classification in the set **C** of all classifications. is probability of a data point being classified as **c** in cluster **w**. Then, normalized mutual information (NMI):

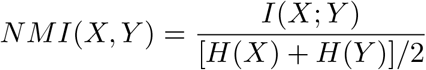

### Benchmark comparison

For all the published datasets we tested, we used the original processed gene expression matrix provided by the authors (Figure. 2a). For data that contained raw read counts, we divided each column by the “total library size” scaling factor to account for differences in sequencing depth, which provide us with reads per million (RPM) value. Log transformations were then applied to the gene count table with a pseudo count of 1.

For each method, we varied the clustering parameters to obtain the optimal clustering results obtainable by each method, as described by the authors in the initial publication and as measured by concordance with the validated results presented in each publication. For t-SNE + k-means, gene filters using median absolute deviation (MAD) or custom filtering criteria were applied. Rtsne (version 0.13) package was used with the default parameters. We used the k reported by the original authors as the input for k-means clustering to identify the specific number of clusters. For SC3 (version 1.5.2), we followed the instructions in the package for gene filtering and clustering. For SINCERA, we used the z-score normalization and automatic cluster identification as described in the original publication. For SEURAT (version 1.4.0.12), we performed t-SNE embedding with the default parameters once and then clustered the data using the DBSCAN algorithm several times, for which we varied the density parameter G in the range of 0.6-3 to find a maximal NMI and reported that number in the summary table.

### Data availability

All melanoma data generated in this study have been deposited in the Gene Expression Omnibus (GEO) database under accession code GEO108397 and SRP127299. All published datasets (Figure 2a) were downloaded from the accession numbers provided in the original publications. These include E-MTAB-3321 (ref Goolam), GSE51372 (ref Ting), GSE45719 (ref Deng), and GSE60361 (ref Zeisel). All analyses were performed using a custom R-based software package available for download at <u>https://github.com/naikai/sake</u>.

## Acknowledgments

YJH is funded through generous supports from the Ministry of Education in Taiwan and The Florence Gould Foundation. MH is a scholar of the Rita Allen Foundation. We also wish to acknowledge a grant from Amazon Web Services to DM, which provided a virtual demo web server for hosting SAKE analyses on the cloud. We would like to thank Meenhard Herlyn and his lab members for their many helpful comments, suggestions, and for the gift of the 451Lu-BR cells. We would also like to acknowledge help from the Bioinformatics Shared Resource at CSHL, which is supported by a grant from the NIH/NCI.

